# Trans-complementation by the RecB nuclease domain of RecBCD enzyme reveals new insight into RecA loading upon χ recognition

**DOI:** 10.1101/2023.05.14.540676

**Authors:** Theetha L. Pavankumar, C. Jason Wong, Yun Ka Wong, Maria Spies, Stephen C. Kowalczykowski

## Abstract

The loading of RecA onto ssDNA by RecBCD is an essential step of RecBCD-mediated homologous recombination. RecBCD facilitates RecA-loading onto ssDNA in a χ-dependent manner via its RecB nuclease domain (RecB^n^). Before recognition of χ, RecB^n^ is sequestered through interactions with RecBCD. It was proposed that upon χ-recognition, RecB^n^ undocks, allowing RecB^n^ to swing out via a contiguous 70 amino acid linker to reveal the RecA-loading surface, and then recruit and load RecA onto ssDNA. We tested this hypothesis by examining the interactions between RecB^n^ (RecB^928–1180^) and truncated RecBCD (RecB^1–927^CD) lacking the nuclease domain. The reconstituted complex of RecB^1–927^CD and RecB^n^ is functional *in vitro* and *in vivo*. Our results indicate that despite being covalently severed from RecB^1–927^CD, RecB^n^ can still load RecA onto ssDNA, establishing that RecB^n^ does not function at the end of its flexible linker. Instead, RecBCD undergoes a χ-induced intramolecular rearrangement to reveal a RecA-loading surface.

## Introduction

The repair of double-stranded DNA (dsDNA) breaks in wild-type *Escherichia coli* requires the functions of RecA and RecBCD proteins ^1, 2^. RecBCD is a heterotrimeric enzyme consisting of RecB, RecC, and RecD subunits ^3, 4^. In its constitutive state, it binds and unwinds from the broken dsDNA ends and simultaneously degrades the 3’-terminated strand at the entry site more extensively than the 5’-terminated strand ^5, 6^. However, the polarity and intensity of nuclease activity is reversed when RecBCD encounters a Chi (χ) sequence (5’-GCTCGTGG-3’) ^7–10^. The stable binding of χ to the RecC subunit of RecBCD is suggested to prevent the χ-containing ssDNA from passing through the nuclease domain and possibly redirecting it through an alternative exit, resulting in preservation of 3’-ended χ-containing ssDNA ^11^. RecBCD then facilitates RecA nucleofilament formation by loading RecA onto the χ-containing ssDNA ^12^. The RecA nucleofilament subsequently searches for homologous DNA to promote DNA pairing ^13^. Subsequent maturation of recombination intermediates ensues to complete the repair process ^2^.

The RecB subunit has two domains with distinct functions: a 100 kDa N-terminal helicase domain (RecB^1–927^; hereafter referred to as RecB^h^) and a 30 kDa C-terminal nuclease domain (RecB^928–1180^; hereafter referred to as RecB^n^), connected by a 70 amino acid (aa) linker region (residues 858 to 927 of RecB) serving as a covalent tether between the two structural domains ^3, 14^. The 70 aa linker region is important for RecBCD activities ^14, 15^. Deletion of RecB^n^ from RecBCD results in a mutant enzyme that fails to load RecA after χ-recognition, suggesting that RecB^n^ plays an important role in RecA-loading ^16^; moreover, RecB^n^ physically interacts with RecA and the areas of RecB^n^ that can interact with the RecA core region were tentatively identified by computational modeling ^17^.

Prior to recognition of χ, RecBCD does not interact with RecA and the surface of RecB^n^ proposed to interact with RecA is buried by interaction with the RecC subunit ^3^. However, several studies indicate that the buried RecA-interacting region of RecB^n^ is revealed upon recognition of χ to facilitate RecA-loading ^16–18^. The discovery that RecB^h^CD has helicase activity but no nuclease actively led to the proposal – the ‘nuclease domain swing model’ (Figure 1a) – that upon χ-recognition, RecB^n^ can swing away via its tether to elicit many of the changes occurring upon χ-recognition ^14^. Another potential consequence of this tethered undocking would be exposure of the RecA-interacting region for subsequent RecA-loading ^4, 15–17^.

**Figure 1.**
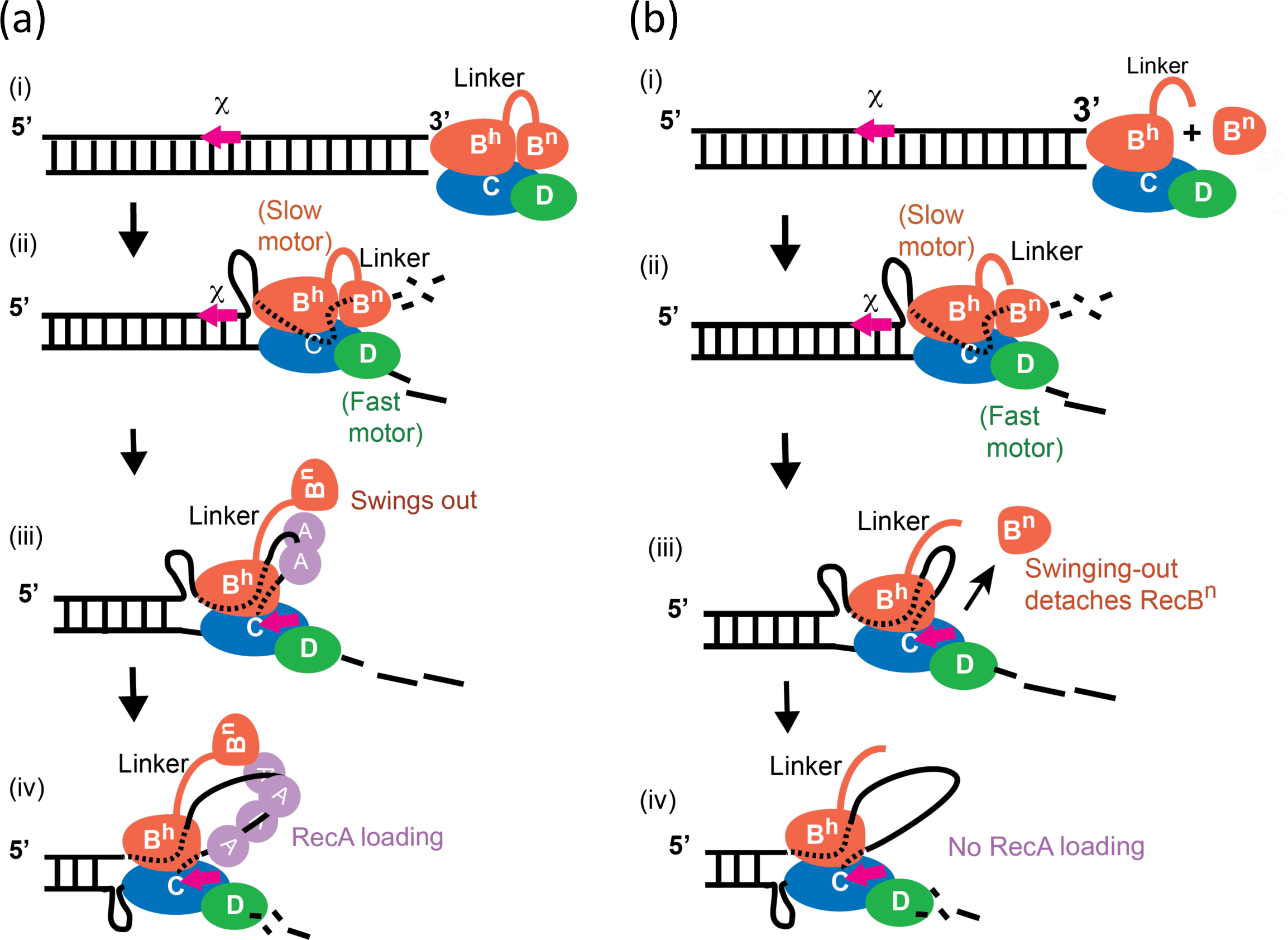
Illustration of the ‘nuclease domain swing model’ for RecBCD. (a) Upon recognition of the χ sequence, the nuclease domain is released to enable the subsequent loading of RecA onto a newly generated χ-containing ssDNA. (i) RecBCD binds to a duplex DNA end. (ii) The duplex DNA is unwound by the slow and fast motors (RecB and RecD respectively) of RecBCD. The nuclease domain preferentially degrades 3’-ended ssDNA-strand over the 5’-ended ssDNA-strand. The RecA-interacting region of the RecB nuclease domain is concealed through the interactions with the RecC subunit. (iii) χ recognition by RecBCD releases the RecB nuclease domain, revealing the RecA-interacting interface of RecB^n^. RecA now interacts with RecB^n^. (iv) RecBCD nucleates RecA onto the χ-containing ssDNA and the RecA-ssDNA filament grows. (b) Illustration of expected outcome of the nuclease swing model for an altered RecBCD with a severed tether. (i) RecB^h^CD and RecB^n^ form a stable complex and the complex binds to a duplex DNA end. (ii) The RecB^h^CD-RecB^n^ complex unwinds and degrades duplex DNA similar to wild-type RecBCD; the RecA-interacting interface of RecB^n^ is concealed. (iii) Upon recognition of χ, the interaction between RecB^h^CD and RecB^n^ is reduced, causing RecB^n^ to detach and swing out from the rest of the enzyme. (iv) As a result, no loading of RecA onto χ-specific ssDNA is observed.

Despite the prominence and value of the nuclease domain swing model, there are no studies showing that RecB^n^ is released upon χ-recognition and that the tether plays the important function of maintaining the RecA-loading domain within proximity of the χ-containing ssDNA. If the model is correct, then a RecBCD enzyme with a severed tether should lose RecB^n^ at χ, and loading of RecA would not ensue (Figure 1b). Here, we show that a functional RecBCD with a severed tether can be reconstituted from a truncated mutant of RecBCD (RecB^1–927^CD) lacking the nuclease domain and a separated RecB nuclease domain (RecB^928–1180^): the severed reconstituted RecBCD complex contains all the residue of the wild-type enzyme but lacks a single peptide bond between amino acid residues 927 and 928. The resulting RecB^h^CD-RecB^n^ complex responds to χ sequences, generates χ-containing ssDNA, and loads RecA onto the χ-containing ssDNA. *In vivo*, RecB^1–927^CD and RecB^928–1180^ complement one another to reconstitute nuclease activity, repair UV-damaged DNA and promote χ-dependent recombination. Based on our results obtained from both genetic and biochemical analyses of RecB^h^CD-RecB^n^ complex, we conclude that covalent tethering is not required for RecBCD functions, implying that RecB^n^ does not function at the free end of a flexible linker. Rather, we conclude that the RecA-interaction site of RecB^n^ is revealed through χ-induced conformational changes within the holoenzyme. We believe that the linker region plays an important role in RecBCD heterotrimer stability and it may serve ensure the association of RecB^n^ with the helicase machinery of RecBCD.

## Results

### RecB^h^CD forms a complex with RecB^n^

We generated and purified a mutant of RecBCD that lacks the nuclease domain of RecB, and we designate it as RecB^h^CD. To test whether the nuclease domain of RecB (RecB^n^) can interact with this truncation of RecBCD, we performed a pull-down assay with 6x His-tagged RecB^n^ (^His^RecB^n^) ^17^ and un-tagged RecB^h^CD. Ni-NTA beads retain protein with the 6x His-tag and, therefore, any un-tagged protein co-eluting with His-tagged protein demonstrates an interaction between the two. As shown in Figure 2a, RecB^h^CD co-elutes with ^His^RecB^n^ from the Ni-NTA beads only in the presence of ^His^RecB^n^. These results show that a complex is formed between RecB^h^CD and ^His^RecB^n^.

**Figure 2.**
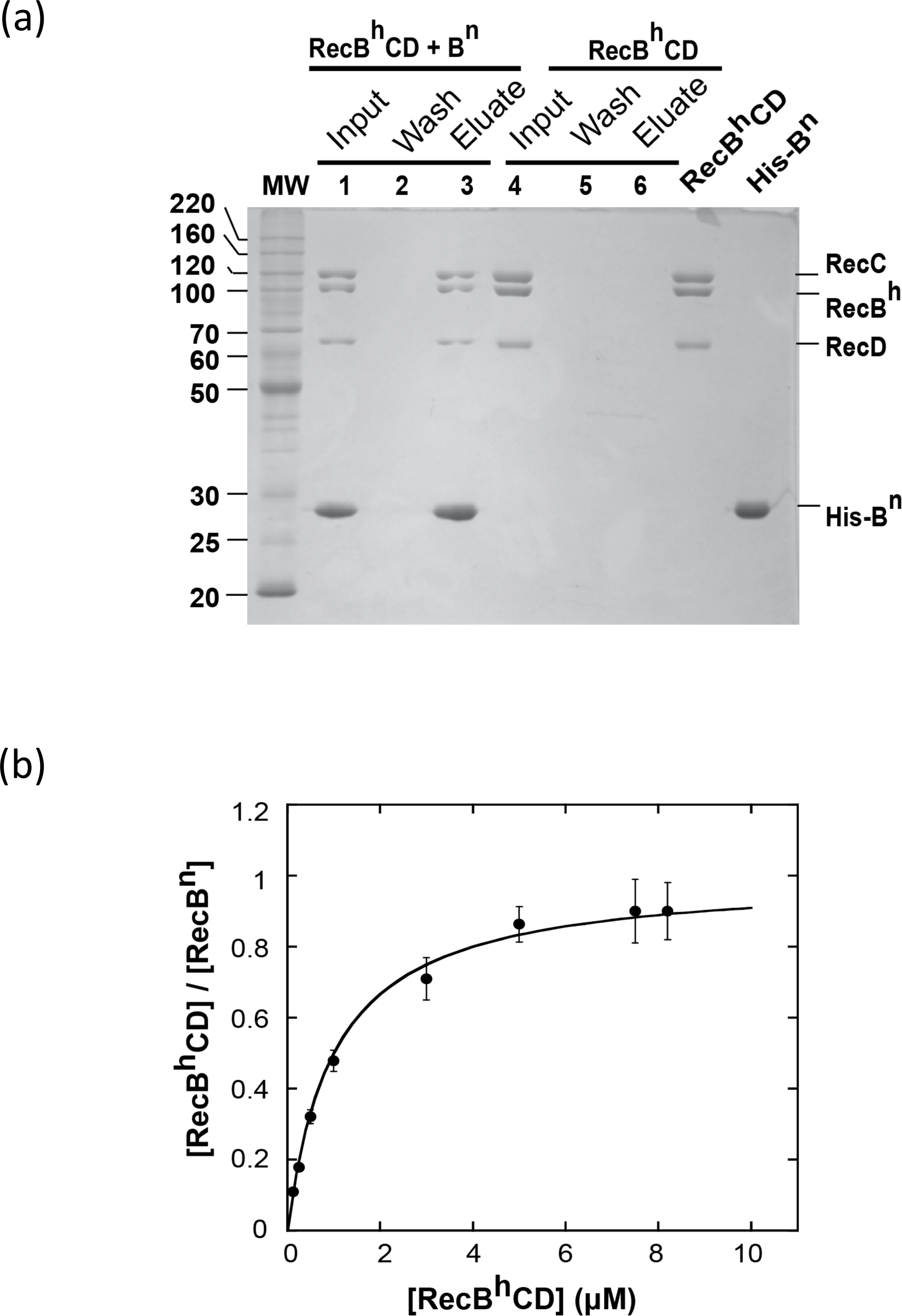
RecB^h^CD forms a stable complex with RecB^n^. The pull-down assay was performed using ^His^RecB^n^ and RecB^h^CD, as described in STAR methods. (a) A representative pull-down experiment; incubation, wash and elution aliquots were analyzed using 12% SDS-PAGE. ^His^RecB^n^ and RecB^h^CD (1 µM each) were incubated with Ni-NTA beads, and an aliquot was removed at the end of incubation and loaded onto Lane 1. Lane 2 is the 3^rd^ wash of the Ni-NTA beads with interaction buffer, and Lane 3 is the elution of Ni-NTA beads with 300 mM imidazole. A control was performed in the absence of ^His^RecB^n^ by incubating Ni-NTA beads with RecB^h^CD (2 µM), and the incubation, wash and elution aliquots were loaded onto Lanes 4, 5, and 6, respectively. The RecB^h^CD and ^His^RecB^n^ markers represent the equivalent of 2 µM and 1 µM, respectively. (b) Determining the binding affinity of RecB^h^CD for RecB^n^. Pull-down assays were performed with 0.5 μM ^His^RecB^n^ and increasing amounts of RecB^h^CD. The ratio of RecB^h^CD to ^His^RecB^n^ eluted from the bead was plotted as a function of total [RecB^h^CD]. The solid line is a simulation using equation (2) and *K*_d_ of 1.1 ± 0.5 μM.

We next performed a pull-down assay with varying the concentrations of RecB^h^CD to measure the affinity between RecB^h^CD and ^His^RecB^n^. The ratio of eluted [RecB^h^CD]/[RecB^n^] is plotted as a function of the total [RecB^h^CD] in Figure 2b, and the data were analyzed with a nonlinear least square (NLLS) algorithm. The analysis yields an apparent dissociation constant (*K*_d_) of 1.1 ± 0.5 μM, and the stoichiometry reaches a plateau value at 0.95 moles RecB^h^CD per mole RecB^n^. The affinity is comparable to the previous observation that RecB^n^ and RecA form a stable complex with an apparent *K*_d_ of ∼1 μM under the same conditions ^17^.

### The RecB^h^CD-RecB^n^ complex produces χ-specific ssDNA fragments

We next examined if this reconstituted complex of RecB^h^CD and RecB^n^ is functional with regards to χ-recognition by studying whether it can generate χ-specific ssDNA fragments ^7^. The formation of χ-specific ssDNA fragments was assayed using a 4.4 kb linear dsDNA containing a triple-χ sequence in each strand and labeled with ^32^P at both 5’-ends (Figure 3a, top panel) ^19^. At a sub-stoichiometric amount of enzyme relative to DNA ends (0.1 nM of enzyme and 2.3 nM DNA ends), different χ-specific ssDNA fragments are produced by wild-type RecBCD depending on which DNA end is the entry site (Figure 3a, top panel). RecB^h^CD alone can only unwind dsDNA to produce full-length ssDNA, and it does not degrade the ssDNA product despite an increased enzyme concentration (Figure 3b). Like RecBCD, the complex formed between RecB^h^CD and RecB^n^ produced χ-specific ssDNA when presented with χ^+^ dsDNA (Figure 3a). Since RecB^n^ has a very low affinity for ssDNA ^20^, it is very unlikely that RecB^n^ is simply binding and digesting the unwound ssDNA products independently of RecB^h^CD. If so, the location of such binding would be random, and therefore, a broad range of degraded ssDNA products should be produced instead of ssDNA products of a specific uniform length. Besides, RecB^n^ alone showed no degradation of linear dsDNA and ssDNA substrates (see last two lanes of Figure 6a).

**Figure 3.**
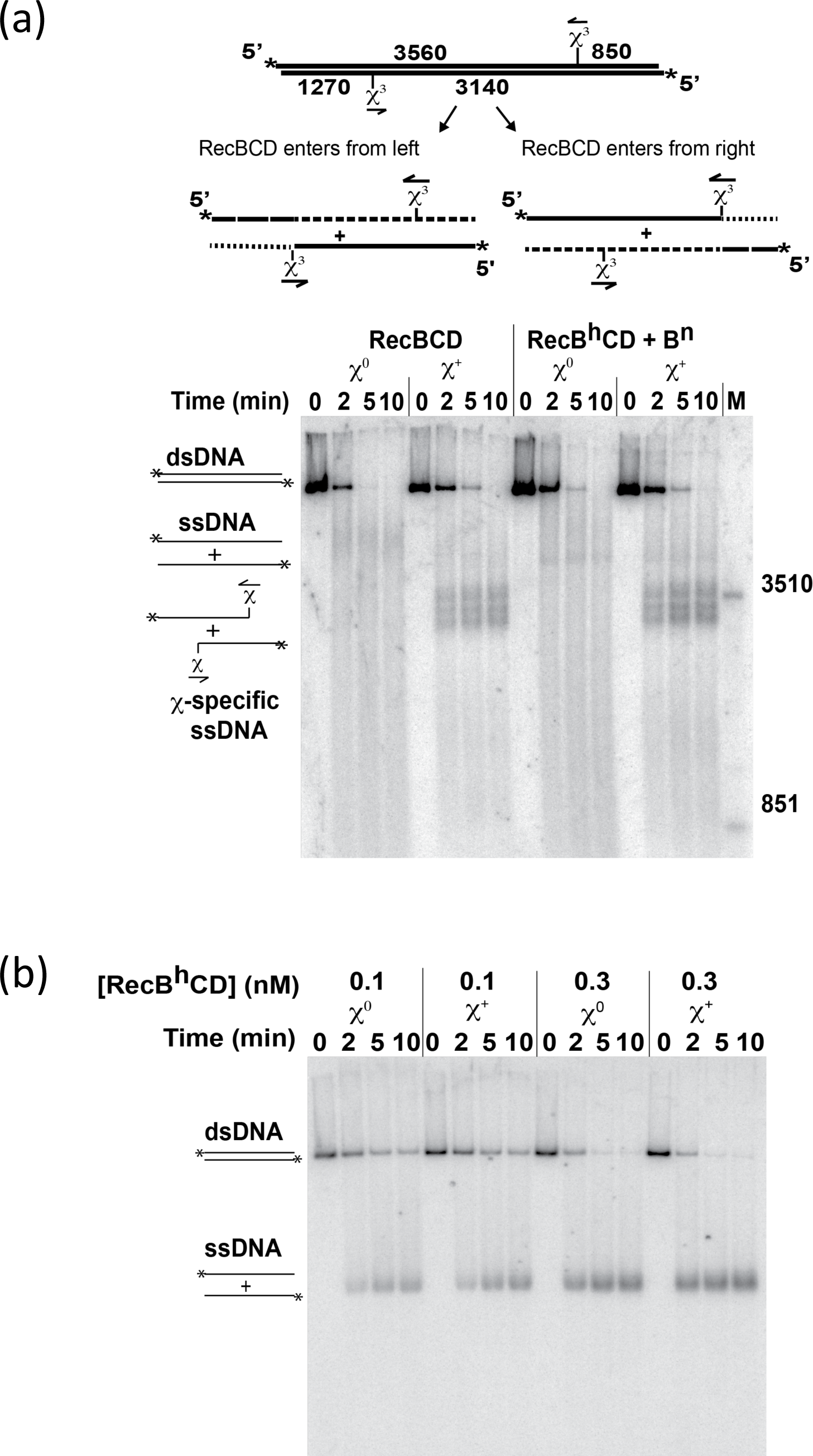
The reconstituted complex of RecB^h^CD and RecB^n^ recognizes χ and produces χ-specific ssDNA fragments. (a) dsDNA (10 μM nt) with χ (χ^+^) or without χ (χ^0^) was reacted with either 0.1 nM RecBCD or 0.1 nM RecB^h^CD plus 1 μM RecB^nuc^. (b) dsDNA (10 μM nt), χ^+^ or χ^0^, was reacted with various concentrations of RecB^h^CD. The location of the triple-χ sequence on the dsDNA is indicated in the inset and the asterisk (*) indicates the position of the γ^32^P label.

### The affinity of RecB^h^CD for RecB^n^ increases in the presence of dsDNA and ATP

We next measured whether the binding affinity of RecB^h^CD for RecB^n^ is affected by the presence of dsDNA and ATP. We measured χ-specific ssDNA production at varying concentrations of RecB^n^ while holding the concentration of RecB^h^CD constant at 0.1 nM. Under the conditions where dsDNA ends are in large excess of RecB^h^CD, all RecB^h^CD as well as RecB^h^CD-RecB^n^ complexes, are bound to the dsDNA ends. Therefore, the fraction of χ-containing fragments generated relative to the total amount of DNA equals to the ratio of RecB^h^CD-RecB^n^ complex to RecB^h^CD bound at the ends of dsDNA. Thus, we can monitor the formation of the RecB^h^CD-RecB^n^ complex as a function of [RecB^n^] and determine the binding affinity. Figure 4a shows a representative experiment. When the fraction of χ-specific ssDNA produced is plotted against total [RecB^n^], a *K*_d_ of 0.4 ± 0.1 μM is obtained from NLLS analyses of the data (Figure 4b). The affinity for RecB^h^CD binding to RecB^n^ is increased by ∼three-fold in the presence of dsDNA and ATP. Importantly, there is good agreement between the pull-down experiment and enzymatic assay.

**Figure 4.**
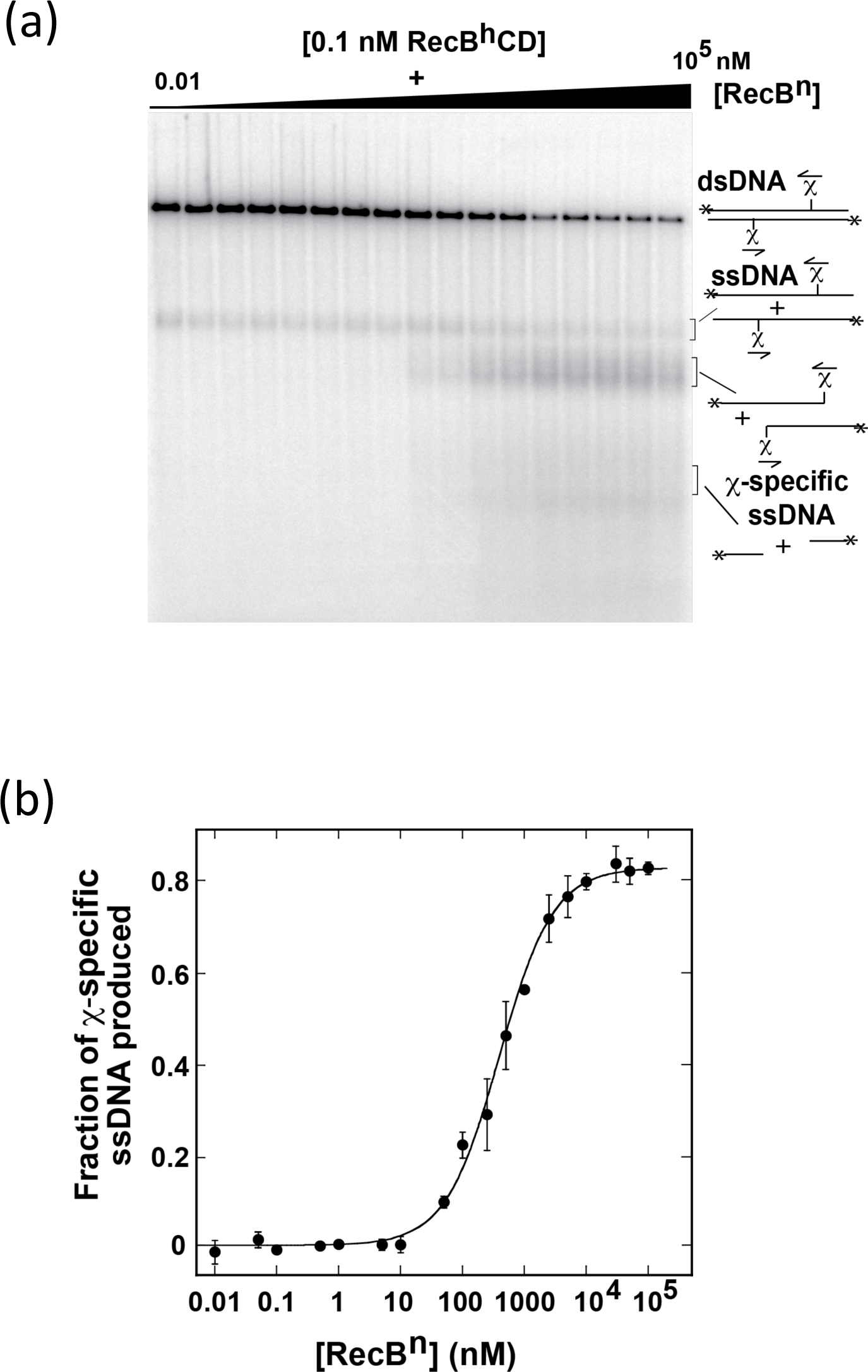
The affinity of RecB^n^ for RecB^h^CD is increased in the presence of dsDNA and ATP. χ-specific ssDNA production was performed with 0.1 nM RecB^h^CD, 20 μM (nt) χ^+^ DNA and varying amounts of RecB^n^. A representative experiment is shown in (a). The fraction of χ-specific ssDNA produced was quantified and plotted as a function of the total [RecB^n^] in (b). Data were analyzed (as described in STAR Methods) and yield an apparent *K*_d_ of 0.4 ± 0.1 μM. The solid line is a simulation using equation (4) and *K*_d_ = 0.4 μM.

### RecB^h^CD recognizes χ despite failure to produce χ-specific fragments

We have shown RecB^h^CD alone cannot produce χ-specific ssDNA but can produce χ-specific ssDNA when it associates with RecB^n^ (Figure 3a and 3b). This suggests that RecB^n^ could play two possible roles within the reconstituted RecB^h^CD-RecB^n^ complex. First, RecB^h^CD alone may not recognize the χ sequence but association with RecB^n^ restores its ability to cleave the DNA up to the 3’-side of χ and then to recognize and attenuate at the χ site. Conversely, it is possible that RecB^h^CD intrinsically recognizes χ, but it cannot produce χ-specific ssDNA due to lack of nuclease activity. To distinguish between these two possibilities, we next examined whether RecB^h^CD can recognize χ or not.

Previous studies have established that the recognition of χ, under conditions of limiting free [Mg^2+^], inhibits DNA unwinding by RecBCD ^21^. The time-course of unwinding reaches an apparent plateau at less than the total amount of dsDNA present. This apparent inhibition is reversed by the addition of excess Mg^2+^, resulting in a burst of unwinding activity, and nearly complete dsDNA unwinding. This results in a distinctive biphasic time-course of unwinding, which is a characteristic behavior of χ-recognition. When χ is absent (χ^0^), neither inhibition at low [Mg^2+^] nor significant biphasic kinetics occur. We performed the same assay to determine whether RecB^h^CD exhibits the same χ-dependent reversible inhibition of dsDNA unwinding. The fraction of DNA unwound was obtained from quantification of gels, and the time courses are plotted in Figure 5. The amount of χ-containing DNA unwound by both RecBCD and RecB^h^CD in 1 mM Mg(OAc)_2_ reaches the same apparent plateau at 40 min. However, upon addition of excess Mg^2+^ to a final concentration of 10 mM, the unwinding rate increased in both cases, indicating that both RecBCD and RecB^h^CD recognize the χ sequence. Thus, RecB^h^CD has the intrinsic capacity to recognize χ, and RecB^n^ has no obligatory role in χ-recognition by the RecB^h^CD-RecB^n^ complex; RecB^n^ however is essential for nuclease activity and the related χ-specific ssDNA formation. This finding is in full agreement with earlier studies for RecBC, lacking RecD, enzyme ^21^.

**Figure 5.**
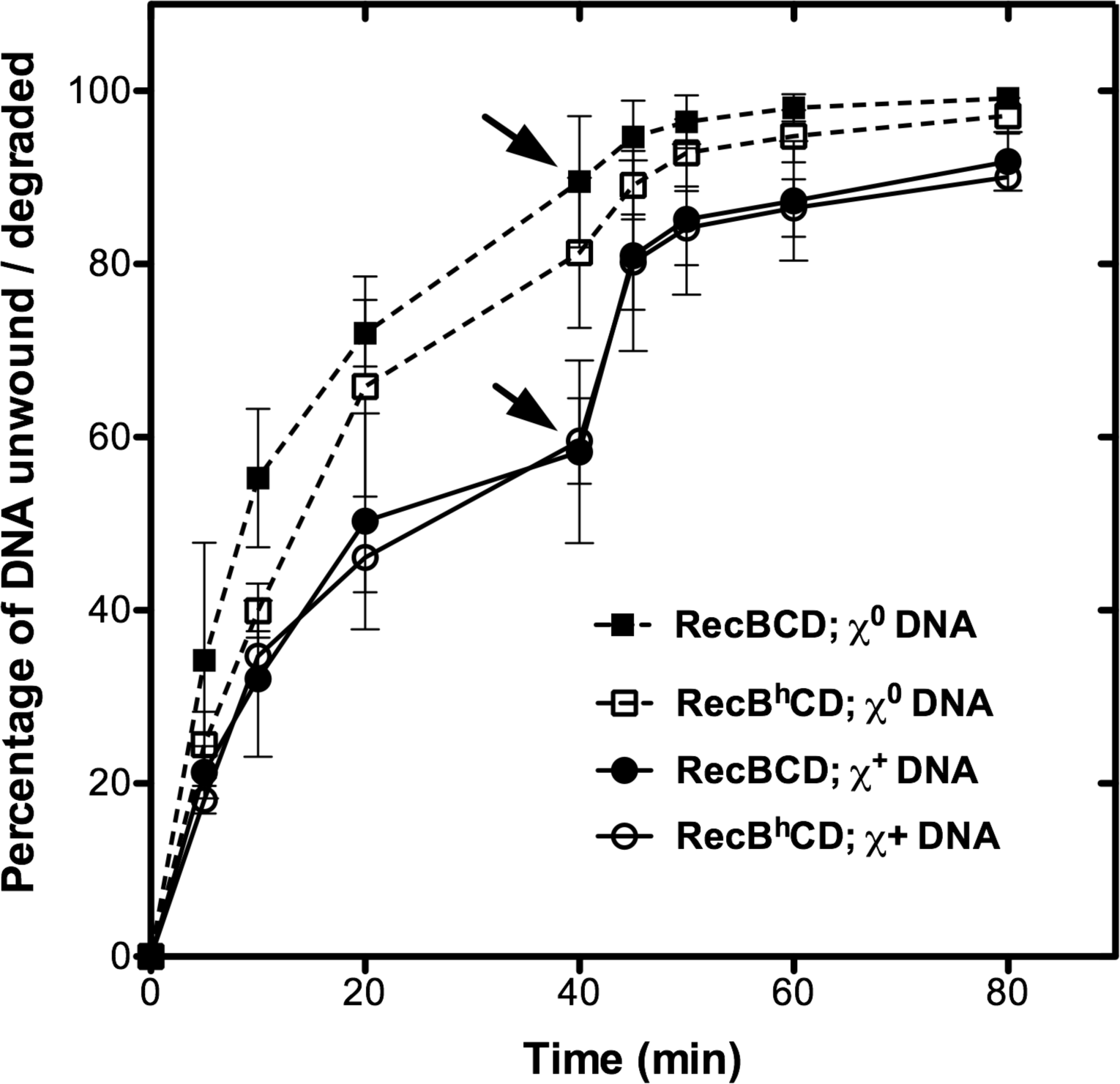
RecB^h^CD recognizes χ as shown by reversible inactivation of dsDNA unwinding upon χ recognition. Either χ^0^ or χ^+^ DNA (10 μM nt) was reacted with 0.1 nM RecBCD or 0.2 nM RecB^h^CD. Aliquots were removed at the times indicated, and excess Mg^2+^ was added after 40 min (indicated by arrows) to a final concentration of 10 mM. Comparison of the time courses of the fraction of DNA unwound by RecBCD with that of RecB^h^CD is shown. Data are the mean ± SEM from two independent experiments.

### The RecB^h^CD-RecB^n^ complex facilitates χ-dependent RecA-loading

RecB^n^ is essential for the RecA-loading process ^16^. So, although the reconstituted RecB^h^CD-RecB^n^ complex produced χ-specific ssDNA fragments, whether it could load RecA onto that ssDNA needed to be determined. Consequently, we measured RecABCD-dependent joint molecule formation by RecBCD, RecB^h^CD, and RecB^h^CD-RecB^n^ enzymes as described previously ^7, 12^. The results show that formation of RecA-dependent, χ-dependent joint molecules (the faster migrating “χ-dep” joint molecules in Figure 6a) by the RecB^h^CD-RecB^n^ complex is similar to that of the wild-type RecBCD enzyme. In contrast, RecB^h^CD alone failed to produce χ-dependent joint molecules and produced only χ-independent joint molecules (from the full-length ssDNA products of unwinding) at a slower rate as previously observed ^7, 12^. The yields of χ-dependent joint molecule formed after 10 minutes were 14.4 ± 1.4% and 13.4 ± 1.1% of the input linear dsDNA substrate for wild-type RecBCD and RecB^h^CD-RecB^n^ enzymes, respectively (Figure 6b). RecB^h^CD alone produced χ-independent joint molecules (4.0 ± 0.4% at 10 minutes); this slower rate is comparable to that of mutant RecBCD enzymes that lack RecA-loading function ^16, 19^. Incubation of RecB^n^ domain alone with linear dsDNA or with ssDNA for 10 min showed no effect (Figure 6a, last two lanes). These findings affirm that the RecB^h^CD-RecB^n^ complex is capable of loading RecA onto the χ-containing ssDNA, and importantly, that the covalent attachment – tethering –between the RecB helicase and RecB^n^ domains is not necessary for the RecA-loading process

**Figure 6.**
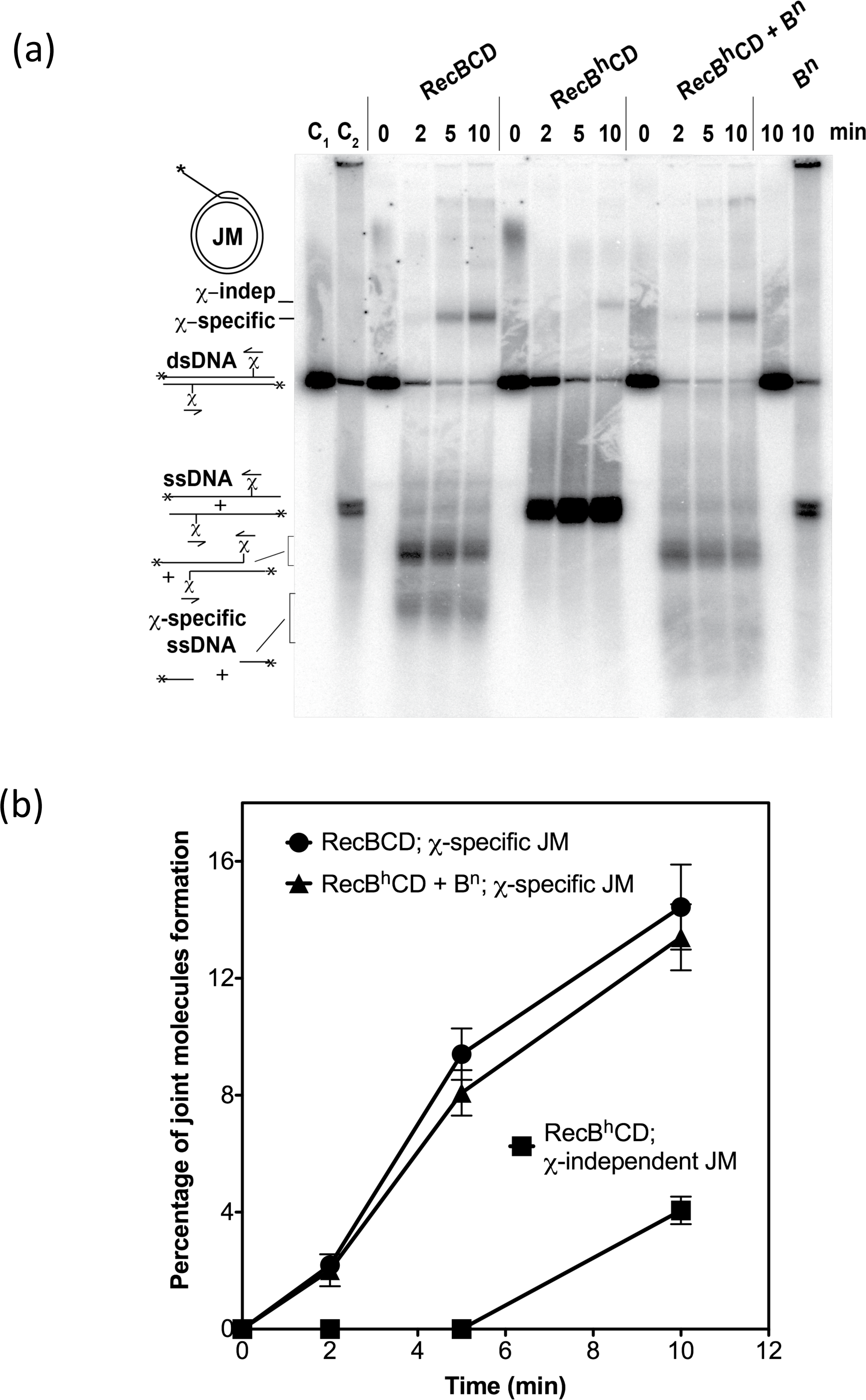
RecB^h^CD-RecB^n^ loads RecA onto χ-containing ssDNA to form χ-specific joint molecule. (a) The χ-dependent (χ-dep) and χ-independent (χ-indep) joint molecules (JM) formed, and the χ-specific ssDNA fragments produced as a function of time. C_1_ and C_2_ are control lanes with 5’-labeled linear dsDNA and ssDNA (heat-denatured dsDNA), respectively. The last two lanes are the reaction mixtures containing linear dsDNA and ssDNA with RecB^n^ alone. (b) Percentage of joint molecules formed relative to the input of linear dsDNA are plotted. Data points are the percentage of joint molecules formed by RecBCD (circles), RecB^h^CD-RecB^n^ (triangles), and RecB^h^CD (rectangles) proteins. Data are the mean ± SEM from three independent experiments.

### The RecB^h^CD-RecB^n^ complex is DNA repair and nuclease proficient *in vivo*

To corroborate our *in vitro* findings, we determined the *in vivo* proficiency of RecB^h^CD-RecB^n^ with regards to its DNA repair function by measuring sensitivity to ultraviolet (UV) irradiation, and to its nuclease activity by measuring infectivity of T4 *2*^−^ phage. Cells expressing RecB^h^CD-RecB^n^, RecB^h^CD, and wild-type RecBCD proteins on plasmids in a *recBCD* null strain were compared. As shown in Figure 7, cells expressing RecB^h^CD protein alone are highly UV sensitive, comparable to the strain that lacks *recBCD* genes (Figure 7). Interestingly, co-expression of RecB^n^ and RecB^h^CD proteins alleviated the UV sensitivity: the ability to repair the UV-induced DNA damage is comparable to the cells that are expressing wild-type RecBCD protein. This result suggests that the RecB nuclease domain is indeed capable of interacting *in trans* with RecB^h^CD enzyme *in vivo* and protecting cells from UV-induced DNA damage.

**Figure 7.**
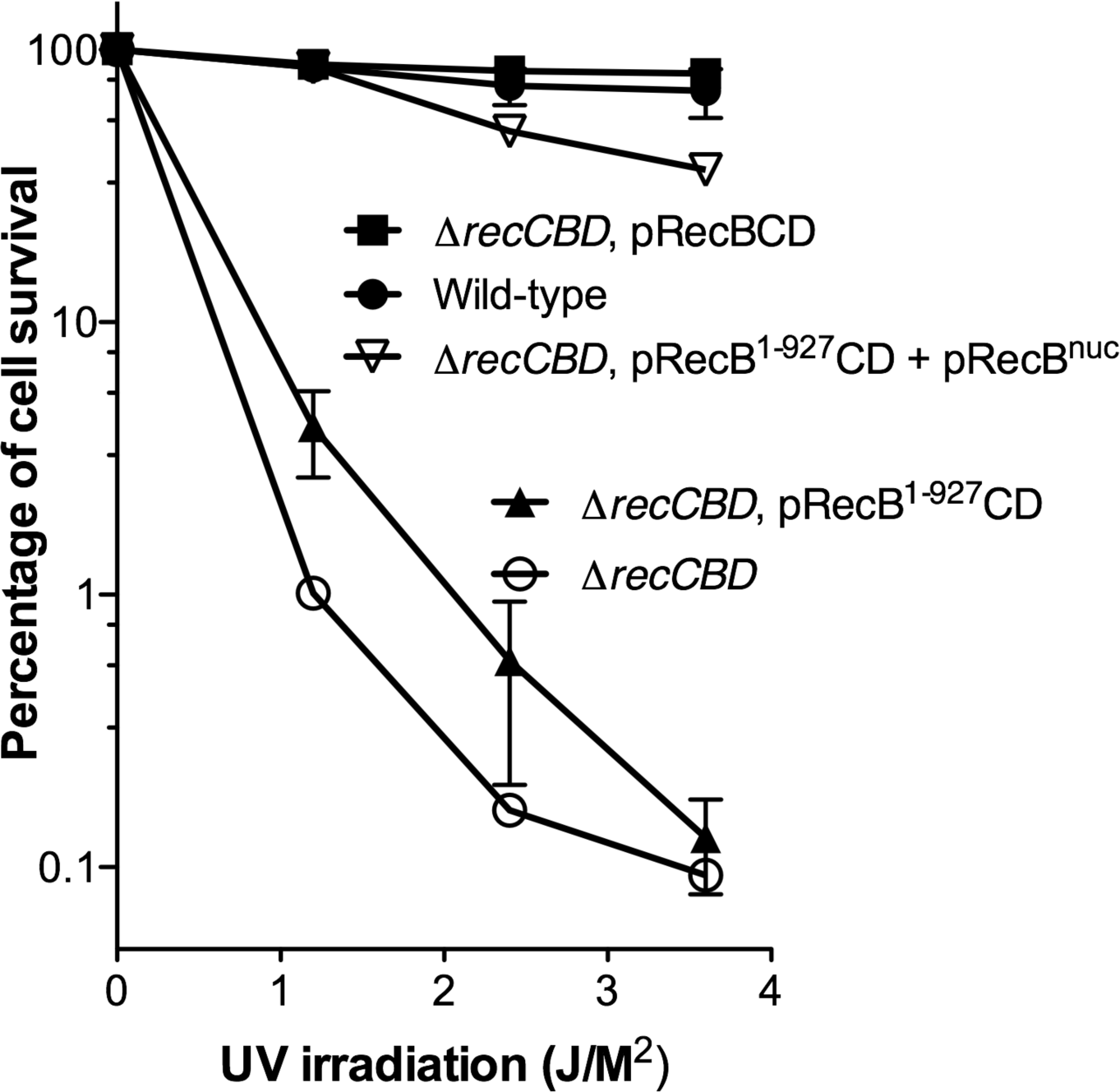
Co-expression of RecB^h^CD and RecB^n^ proteins confers DNA repair proficiency to *E. coli* cells. A *recBCD* null strain expressing wild-type RecBCD, RecB^h^CD, RecB^h^CD-RecB^n^ proteins was subjected to UV irradiation. Percentage of cells surviving at various doses of UV irradiation are compared. Error bars of some data-points are minimal, hence not visible on the graph. Data are the mean ± SEM from the three independent experiments.

If the RecB nuclease domain interacts with RecB^h^CD *in vivo*, then it is expected to have nucleolytic activity. To determine whether the RecB^h^CD-RecB^n^ complex has nucleolytic activity, we assayed T4 *2*^−^ phage sensitivity. Wild-type T4 phage with a functional *gene 2* product is resistant to RecBCD-dependent DNA degradation due to the protection afforded by the binding of gene 2 protein to dsDNA ends of T4 phage DNA. However, T4 *2*^−^ phages lacking gene 2 product make plaques only on the strain lacking RecBCD nuclease activity. Table 1 shows plaque formation by T4 *2*^−^ in different backgrounds. In the strain expressing RecB^h^CD and RecB^n^ proteins, viability T4 *2*^−^ phage is decreased by almost a thousand-fold compared to the strain expressing RecB^h^CD alone and is reduced only ∼10-fold compared to wild-type RecBCD. These results further confirm the ability of RecB^n^ to associate *in trans* with RecB^h^CD *in vivo* to confer exonucleolytic activity to the cells.

**Table 1:** Exonuclease activity of RecBCD mutants *in vivo*.

| *E. coli* strain | T4 phage titer<sup>a</sup> | T4 2<sup>-</sup> phage titer<sup>a</sup> |
| --- | --- | --- |
| AB1157 (WT) | 0.8 ± 0.1 | 1.8 ± 0.1 × 10<sup>-4</sup> |
| $\Delta recCBD::FRT$ | 1 | 1 |
| $\Delta recCBD::FRT$ (pRecBCD) | 1.0 ± 0.1 | 2.5 ± 0.3 × 10<sup>-4</sup> |
| $\Delta recCBD::FRT$ (pRecB<sup>1-927</sup>CD) | 2.0 ± 0.1 | 1.1 ± 0.1 |
| $\Delta recCBD::FRT$ (pRecB<sup>1-927</sup>CD & pRecB<sup>n</sup>) | 0.8 ± 0.2 | 2.9 ± 1.7 × 10<sup>-3</sup> |

<sup>a</sup>Relative values: T4 wild-type or T4 2<sup>-</sup> phage titers from the indicated strains are divided by the titer on $\Delta recCBD::FRT$ strain. Data are the mean ± SEM from three independent experiments.

### The RecB^h^CD-RecB^n^ complex promotes χ-stimulated homologous recombination ***in vivo*.**

We further investigated the functionality of the RecB^h^CD-RecB^n^ complex in *vivo* by measuring χ-stimulated recombination between *λ red gam* phages which lack both the RecBCD-inhibiting Gam protein, and the λ recombination system, Red ^22^. Cells expressing RecB^h^CD-RecB^n^ complex showed a ∼10-fold increase in *λ* recombinants and a ∼4-fold higher χ-stimulated crossover activity compared to the cells expressing RecB^h^CD alone (Table 2); these values approach those of wild-type RecBCD. These results further confirm that the RecB^n^ indeed interacts with RecB^h^CD *in vivo* to reconstitute a holoenzyme that is functional for χ-stimulated recombination in cells.

**Table 2:** Recombination proficiency and **χ** activity of the RecBCD mutants *in vivo*.

| Strains | $\lambda$ Recombinant frequency<br>(J<sup>+</sup> R<sup>+</sup> recombinants, %) | | $\chi$ activity |
| --- | --- | --- | --- |
|  | Cross 1 | Cross 2 |  |
|  | (1081 × 1082) | (1083 × 1084) |  |
| AB1157 (WT) | 8.7 ± 0.6 | 7.7 ± 0.6 | 5.1 ± 0.1 |
| $\Delta recCBD::FRT$ | 0.1 ± 0.01 | 0.2 ± 0.05 | 1.1 ± 0.1 |
| $\Delta recCBD::FRT$ (pRecBCD) | 8.7 ± 0.01 | 7.9 ± 0.3 | 5.7 ± 0.9 |
| $\Delta recCBD::FRT$ (pRecB<sup>1-927</sup>CD) | 0.9 ± 0.2 | 0.7 ± 0.4 | 1.5 ± 0.2 |
| $\Delta recCBD::FRT$ (pRecB<sup>1-927</sup>CD & pRecB<sup>n</sup>) | 7.2 ± 0.7 | 6.9 ± 0.04 | 4.1 ± 0.3 |

Data are the mean ± SEM from two independent experiments.

## Discussion

RecBCD enzyme plays an essential role in the recombinational DNA repair of dsDNA breaks in *E. coli*. It is a multifunctional enzyme that initiates the DSB repair by nucleolytically processing the DSB. Its DNA resection activity serves as a prototype for dsDNA processing and mediator function in all organisms. Homologs or analogs specifically involved in resection exist in all organisms: in Eukaryotes, the Sgs1-Dna2-RPA-Top3-Rmi1 complex in *Saccharomyces cerevisiae* and the BLM-DNA2-RPA-TOP3α-RMI-RMI2 complex in humans comprise functional analogs of the χ-activated RecBCD complex ^2, 23–25^.

RecBCD also possesses another essential activity: it mediates the delivery of RecA onto ssDNA that is complexed with SSB. This loading of RecA by RecBCD in response to χ-activation is a crucial step of recombinational dsDNA break repair; however, the detailed mechanism by which this occurs is not fully understood. RecBCD enzyme, specifically the nuclease/RecA-loading domain, is known to interact with RecA and to load RecA onto ssDNA, but only after recognition of χ. The isolated nuclease domain of RecB interacts directly with RecA and regions potentially responsible for such interactions were identified ^17^. Before recognition of χ, the RecA-interacting surface on RecB^n^ is occupied through the interactions with the RecC subunit ^3^. In response to χ, by an unspecified structural mechanism, the concealed RecA-interacting interface of RecB^n^ is revealed to facilitate RecA-loading onto ssDNA ^12, 16, 17, 26^.

One model, the ‘nuclease domain swing model’, described in the Introduction, proposed that upon χ-recognition, conformational changes in RecBCD liberate RecB^n^ from its binding site on RecC resulting in the exposure of the RecA-interacting surface of RecB^n^. The 70 aa linker was envisioned to tether the loading-domain in proximity to the newly generated ssDNA (Figure 1a-(iii)). RecA then interacts with RecB^n^ and is nucleated on the χ-containing ssDNA (Figure 1a-(iv)) ^4, 14–17^. A prediction of the simplest form of this model is that without a tether, if RecB^n^ were to be released, none of the RecB^n^-dependent events downstream of χ-recognition would occur.

We examined this model by severing the link between RecB^h^CD and RecB^n^. We discovered that we could reconstitute the activities of wild-type RecBCD, despite the absence of a contiguous covalent linker. Pull-down experiments showed that the RecB^h^CD forms 1:1 complex with RecB^n^ in the absence of DNA with an affinity, similar to that observed between RecA and RecB^n^ (*K*_d_ ∼ 1 μM) ^17^. In the presence of dsDNA and ATP, the affinity between RecB^h^CD and RecB^n^ increases slightly (*K*_d_ = 0.4 ± 0.1 μM). As is evident from the enzymatic assays and consistent with the equilibrium binding constant, RecB^h^CD is predominantly free and not bound to RecB^n^ until concentrations greater than ∼0.4 μM (e.g., see Figure 4), suggesting that a function of the linker is to ensure interaction of nuclease and RecA-loading domain with the RecBCD holoenzyme by creating a unimolecular species of RecB^h^ and RecB^n^.

Unexpectedly, we discovered that the reconstituted complex formed between RecB^h^CD and RecB^n^ can process χ-containing dsDNA to produce χ-specific ssDNA as does wild-type RecBCD enzyme. Also, the reconstituted RecB^h^CD-RecB^n^ complex is RecA-loading proficient like the wild-type RecBCD enzyme. Hence, the interactions between RecB^n^ and RecB^h^CD are functional and sufficient for producing χ-specific ssDNA and RecA-loading. This conclusion is supported by *in vivo* observations of decreased plaque formation of T4 *2*^−^ phage in RecB^h^CD-RecB^n^ expressing strain, showing that RecB^n^ and RecB^h^CD form an active nuclease complex in cells. Furthermore, the RecB^h^CD-RecB^n^ complex showed χ-dependent crossover hotpot activity in classical lambda phage crosses, confirming reconstitution of a functional RecBCD complex. These physiological measurements are slightly reduced compared to wild-type RecBCD, implying incomplete saturation of RecB^h^CD with RecB^n^, again consistent with a function of the tether being to produce a single polypeptide comprising RecB^h^CD and RecB^n^ rather than relying on bimolecular complex formation.

However, our findings that RecB^h^CD and RecB^n^ can reconstitute typical RecBCD functions is in contradiction with the swing-model for RecA loading. Specifically, the results show that an intact covalent tether is unnecessary; thus, RecBCD has reorganized itself after χ-recognition and the RecA interaction site of RecB^n^ was revealed without covalent tethering. Thus, in contrast to the prevailing model, non-covalent interactions between RecB^n^ and RecB^h^CD play an important role in the association of RecB^n^ with the rest of the enzyme even after recognition of χ. Based on our results, we propose that χ recognition causes conformational changes in both the RecB^n^ and RecB^h^CD proteins. As a result, rather than swinging freely as a ball on a tether, RecB^n^ functions while remaining attached to the rest of the enzyme but in a different conformation. These unexpected findings raise the question of what is the function of the 70-aa linker? Our attempts to purify truncated RecB proteins with shorter linkers (RecB^1–, 866^ and RecB^1–898^) revealed that the RecBCD heterotrimer was unstable, suggesting that the linker is likely a “strap” that is important to the stability of the RecBC complex within the heterotrimer.

Consequently, we propose a new model of RecA-loading by RecBCD in response to χ. RecBCD unwinds and degrades the duplex DNA from the DNA end (Figure 8(i)). Before the recognition of χ, the RecB^n^ remains associated with the RecC subunit, and the RecA-loading surface is concealed (Figure 8(ii)). RecBCD recognizes χ and the recognition of χ induces conformational changes in RecBCD causing rearrangement of interactions with the RecA-loading domain, RecB^n^ (Figure 8(iii)). These structural rearrangements both reveal the RecA-interaction interface and modify the degradative focus of RecB^n^ to produce 3’-ended, χ-containing ssDNA and facilitate RecA-loading on to it (Figure 8-(iv)).

**Figure 8.**
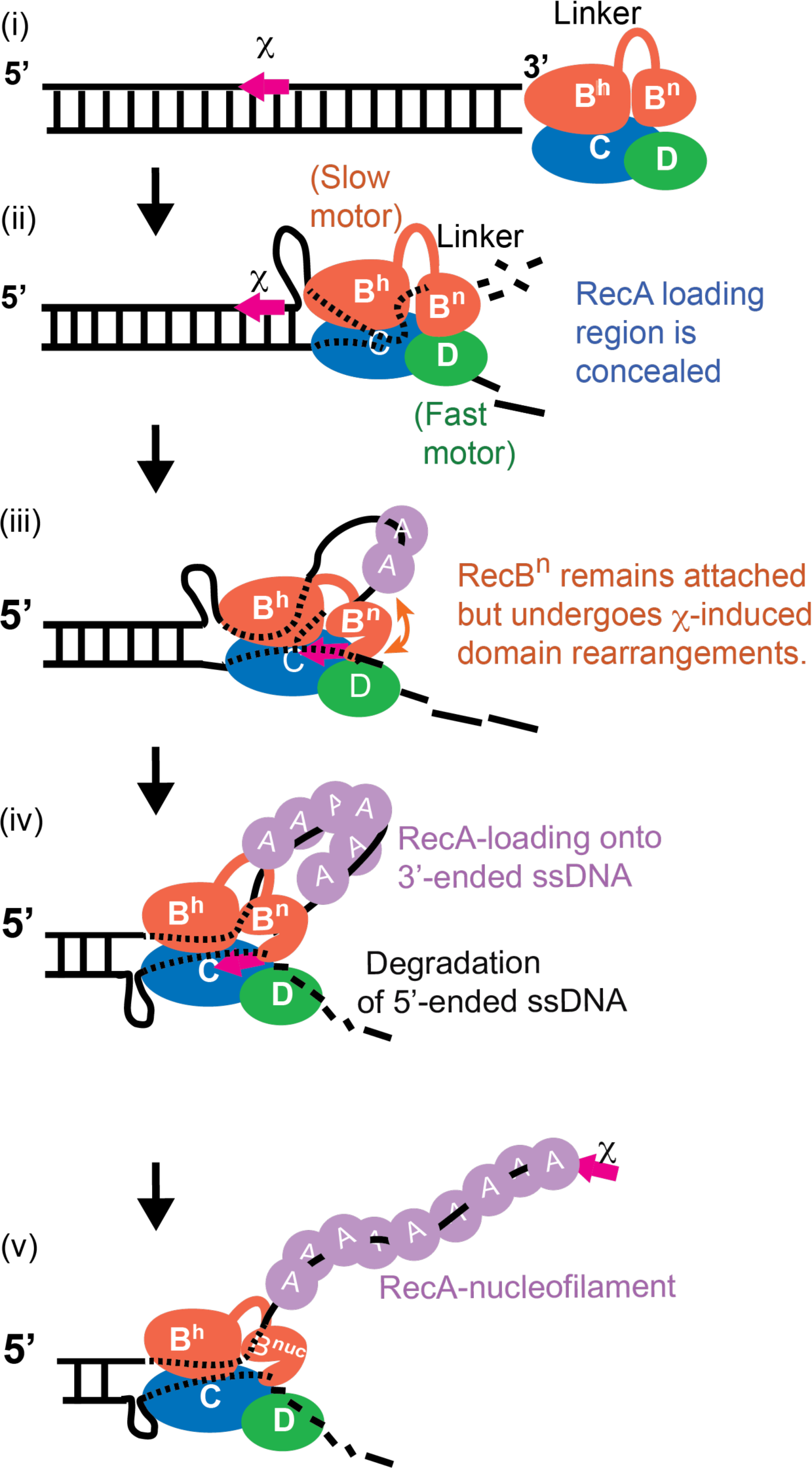
Model illustrating the prospective mechanism of χ-dependent RecA-loading by the RecBCD enzyme and double-strand break repair. (i) RecBCD binds to a duplex DNA end. (ii) The slow motor (RecB) and the fast motor (RecD) unwind duplex DNA. The differential translocation speed between two motor creates a loop ahead of slow RecB motor. The unwound duplex DNA is degraded by RecB nuclease domain, and the 3’-end ssDNA is preferentially degraded over the 5’-ended ssDNA. (iii) RecBCD recognizes the χ sequence, and χ remains transiently bound to the RecC subunit. The χ-induced conformational changes in RecBCD leads to domain rearrangements of RecB nuclease while remaining attached to the rest of the RecBCD enzyme. (iv) The structural rearrangements affecting RecB^n^ cause attenuation of nuclease activity leaving χ-containing ssDNA with the χ sequence at the 3’-end and reveal the RecA-interacting interface to load RecA onto the χ-containing ssDNA. (v) After RecA nucleation, growth of the RecA filament needed for the homology search ensues.

## Acknowledgements

We would like to thank Drs. Gerald Smith and Susan Amundsen of Fred Hutchinson Cancer Research Center, Seattle, USA for generously providing the strains and phages for the *in vivo* nuclease and recombination assays. These studies were supported by NIH grant R35 GM131900 to S.C.K.

## Authors contributions

S.C.K., and M.S. conceived the study. S.C.K., M.S., J.W., and T.L.P. designed experiments. Experiments were performed by J.W., and T.L.P. and Y.K.W. assisted T.L.P. in genetic experiments. T.L.P., J.W., and S.C.K. analyzed the data and wrote the manuscript. S.C.K. acquired funding and supervised the project.

## Inclusion and diversity statement

We support inclusive, diverse, and equitable conduct of research.

## STAR Methods

### Key resources table

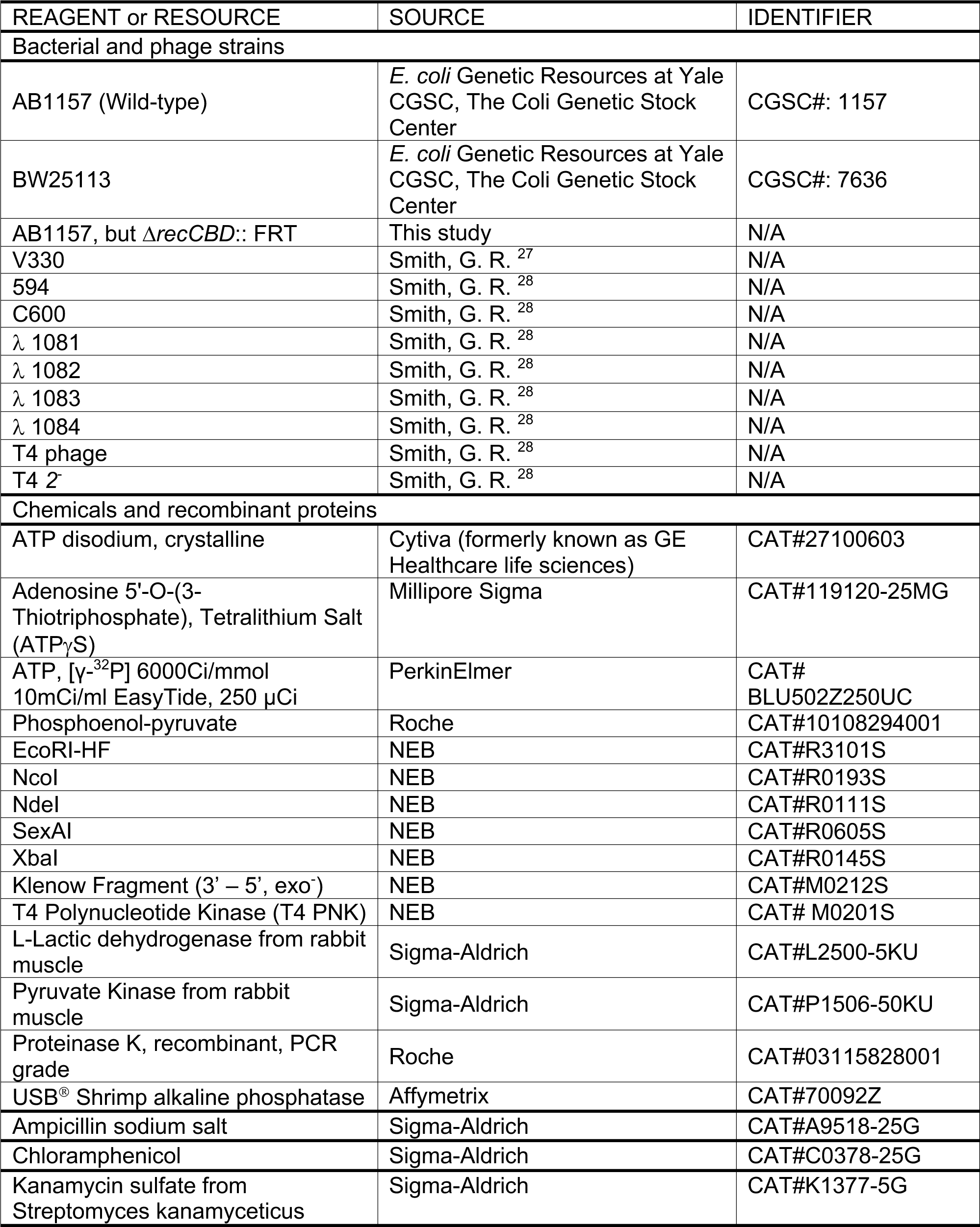
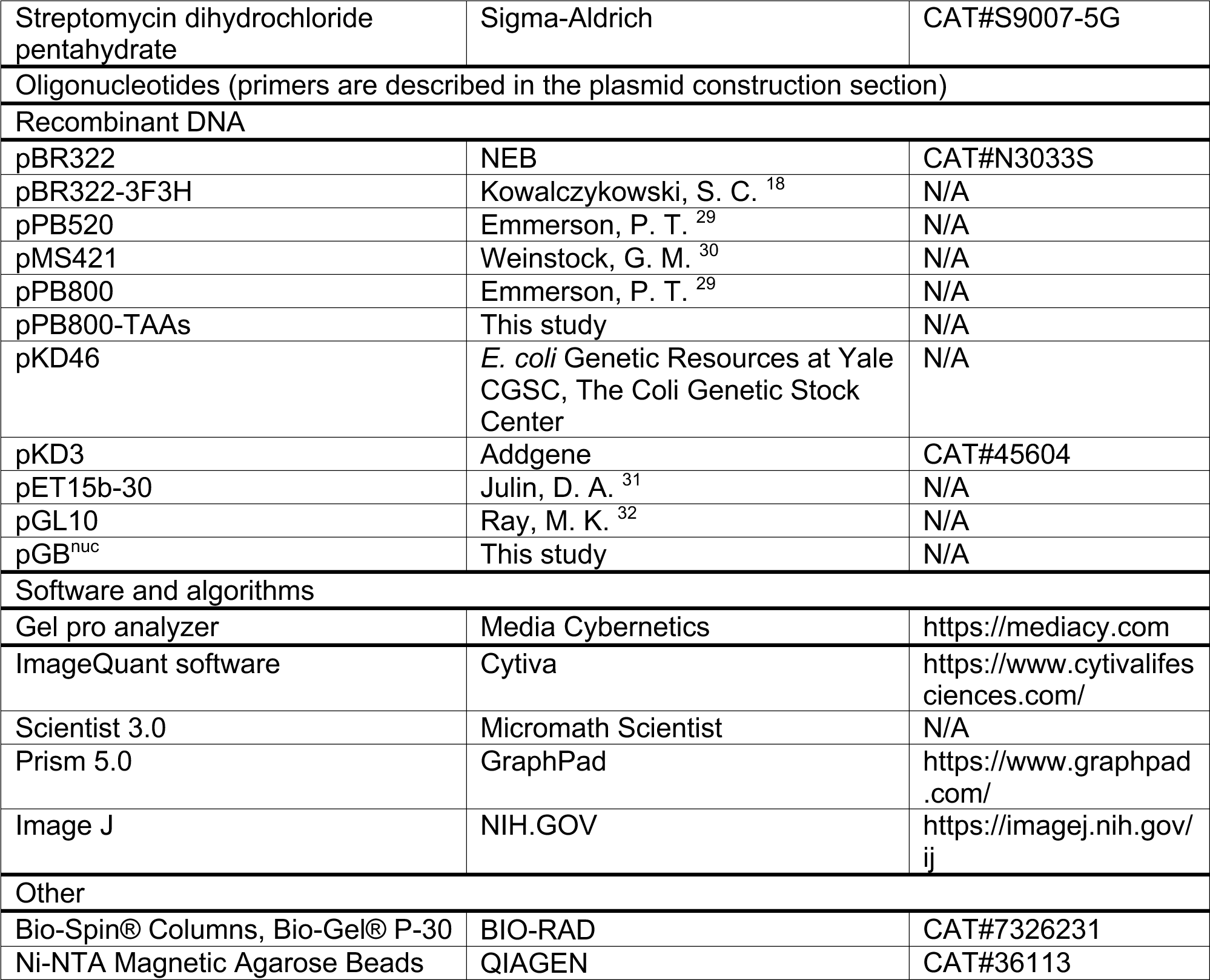

### Lead contact

Further information and requests for resources and reagents should be directed to and will be fulfilled by the Lead contact, Stephen C. Kowalczykowski.

### Materials availability

No restrictions for use of materials from this study.

### Experimental model and subject details

#### Bacterial strains and phages

The following *Escherichia coli* strains were used in the experiments as described in the method details. AB1157 cells as wildtype, the *recCBD* operon deleted AB1157 (1−*recCBD*::*FRT)*, V330 (1− *(recC-argA) 234 λ^− F−^)* ^27^, 594 (*lac-3350 galK2 galT22 rpsL179 λ^− F−^)*, and C600 (*thr-1 leuB6 thi-1 lacY1 tonA21 supE44 rfbD1 λ^− F−^*) ^28^. The following phage strains were also used in the study. λ 1081 (*susJ6 b1453 cI857 χ^+^D123*), λ 1082 (*b1453 χ^+^D123 susR5*), λ 1083 (*susJ6 b1453 χ^+^76 cI857*), λ 1084 (*b1453 χ^+^76 susR5*), T4 phage (*gene 2^+^*), and T4 *2*^−^ (*gene 2 amN51*). All the phages were generous gifts from Gerald R. Smith ^28^.

#### Buffers and Reagents

Buffers were made from reagent grade chemicals with Milli-Q water. ATP (Cytiva, USA) and ATPγS (Millipore Sigma, USA) stocks were adjusted to pH 7.5, and concentration was determined spectrophotometrically using ϵ_260_ = 1.54 χ 10^4^ M^−1^ cm^−1^.

### Method details

#### Purification of proteins

6x His-tagged RecB^n^ (^His^RecB^n^) was purified as described ^17^. The plasmid for overexpression of RecB^1–927^ and RecD was made by replacing bases 960 to 965 of pPB800 ^29^ with two TAA stop codons, and this plasmid is then transformed into a *1−recCBD E. coli* strain V330 ^27^ carrying pPB520 (*recC*^+^) ^29^ and pMS421 (*lac*I^q^) ^30^. This strain was grown, lysed, and RecB^h^CD was purified chromatographically in the same way as wild-type RecBCD as described previously ^33^. We also generated two longer truncations of the RecB nuclease domain to generate RecB^1–866^ and RecB^1–898^. We expressed them together with RecC and RecD subunits as was done for the RecB^1–927^ truncation. However, we failed to purify stable heterotrimeric RecBCD complexes with these longer deletions, despite clear expression of all the subunits.

Both RecB^h^CD and ^His^RecB^n^ are stored in buffer B (20 mM Tris-HCl, pH 7.5, 0.1 mM EDTA, 0.1 mM DTT, 100 mM NaCl, 50% (v/v) glycerol) at −80°C. RecB^h^CD and ^His^RecB^n^ have extinction coefficients of ϵ_280_ = 4.2 χ 10^5^ M^−1^ cm^−1^ and 4.6 χ 10^4^ M^−1^ cm^−1^ in buffer B, respectively. These values are determined by comparing the absorbance spectra of aliquots of protein in buffer B to spectra taken in 6 M guanidinium-HCl. The extinction coefficients of either RecB^h^CD or ^His^RecB^n^ in 6 M guanidinium-HCl at 280 nm was calculated from the amino acid sequence as described ^34^. RecA and SSB were purified as described ^35^.

### Preparation of DNA substrates

DNA substrate χ^0^ was prepared from circular plasmid pBR322, while χ^+^ DNA was prepared from pBR322 3F3H, (a pBR322 derivative with two sets of three tandem χ-sequences) ^19^. These circular plasmid DNA were purified by cesium chloride density gradient centrifugation ^36^. The molar concentration of DNA was determined using an extinction coefficient of 6290 M^−1^ cm^−1^ nt^−1^ at 260 nm. Plasmid DNA was linearized using NdeI (NEB, Ipswish, MA) and radioactively labeled at the 5’-end by sequential reaction with shrimp alkaline phosphatase (USB Corp, Cleveland, OH) and T4 polynucleotide kinase (NEB, Ipswish, MA) and [γ-^32^P] ATP (Perkin Elmer, Wellesley, MA) followed by purification with P30 BioSpin columns (Bio-Rad, Hercules, CA). 3’-labeling was performed by adding Klenow fragment (exo^−^) of Polymerase I (NEB, Ipswish, MA) and [α-^32^P] ATP (Perkin Elmer, Hopkinton, MA) to the linearized DNA and purified as described above.

### Generation of 1−*recCBD* strain of *E. coli* and plasmid constructions

#### Generation of 1−*rec*CB*D*::FRT (flipase recognition target (FRT)) *AB1157* strain

For the generation of 1−*recCBD::*FRT AB1157 strain ^37^, the homologous regions of *recC* and *recD* genes (underlined), along with the chloramphenicol acetyltransferase gene (*cat*) of pKD3 plasmid was PCR amplified using the forward primer, 5’-<u>GCTGGTGGAATATACCCATCAACTCGGGCAACCGCGCTGGCACCGCGCCAATCTC</u> GTGTAGGCTGGAGCTGCTTC-3’ and reverse primer 5’-<u>GAGCTTCCTCAAGCATCGCAATATAATCTTCGCCGCTCTGTAAAAGCCGTTTTTCG</u>C ATATGAATATCCTCCTTAG-3’. The purified PCR product was then transformed into a BW25113 strain carrying pKD46 plasmid ^37^. The P1 lysate of BW25113 transformants were later transduced into AB1157 cells. The *cat* gene was removed by FLP recombinase to obtain AB1157 1−*recCBD::*FRT strain as described by Daysenko and Wanner method of one-step inactivation of genes in *E. coli* ^37^.

#### Construction of pPB800-TAAs

A 1.2 kb region of *recB* gene was PCR amplified using a forward primer 5’-CGCTGGCAACGCCTTATTAATCGACATCCAGCCG-3’ and a reverse primer 5’-CGGCTGGATGTCGATTAATAAGGCGTTGCCAGCG-3’ containing two stop codons at the 928^th^ and 929^th^ positions of *recB* gene. The PCR amplified region was digested with NcoI and SexAI enzymes and cloned into the NcoI and SexAI digested pPB800 vector (pKK223-3 based *recB* and *recD* overexpression plasmid ^29^) to obtain pPB800-TAAs plasmid. The pPB800-TAAs contains two TAA stop codons in the places of 928^th^ and 929^th^ amino acids of RecB. It expresses the RecB^1–927^ and RecD polypeptides without RecB^n^ domain. We also constructed vectors for expressing RecB^1–866^ and RecB^1–898^ truncated RecB proteins using following primers. RecB-866 forward primer – 5’-GTTTGCCAGGCAATTTATTAATCGCATAACGCTTC-3’ and RecB-866 reverse primer – 5’-GAAGCGTTATGCGATTAATAAATTGCCTGGCAAAC-3’; RecB-898 forward primer-5’-CAACGATTGCCCGGCTAATGATGGCGCGTCACCAGC-3’ and RecB-898 reverse primer 5’-GCTGGTGACGCGCCATCATTAGCCGGGCAATCGTTG-3’. The DNA sequencing was performed to confirm the integrity of the DNA sequence and presence of stop codons.

#### Construction of pGB^nuc^

The XbaI-EcoRI restriction digested 1.2 kb DNA fragment containing RecB^n^ gene from the pET-15b-30 vector ^31^ was cloned into a low copy number plasmid pGL10 vector ^32^ at the XbaI-EcoRI sites. DNA sequencing was performed to confirm the integrity and presence of RecB^n^ gene, and expression of RecB^n^ was also confirmed.

### Protein-protein interaction assays using Ni-NTA magnetic beads

Ni-NTA magnetic beads (QIAGEN, Valencia, CA) were used to study the interactions between ^His^RecB^n^ and RecB^h^CD as described ^17^. RecB^n^ (0.5 μM) was incubated with various amounts of RecB^h^CD at the indicated conditions in 50 mM Tris-acetate pH 7.5, 50 mM NaCl, 50 mM imidazole, 0.2% Triton X-100 for 15 min. Ni-NTA beads were added to a final concentration of 1%, and the beads were immobilized, washed, and eluted as described ^17^. Eluted fractions were analyzed on 12% SDS-PAGE, followed by staining with Coomassie brilliant blue and quantified on a Gel Pro Analyzer (Media Cybernetics, Bethesda, MD). The amount of protein eluted was calculated by comparison to known quantities of each protein. The binding isotherm was analyzed by assuming one RecB^n^ binds to one RecB^h^CD as described in Scheme 1:

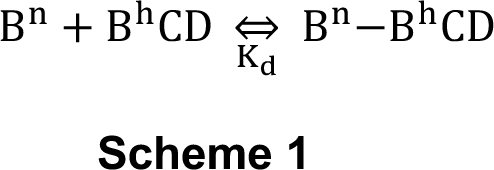

where B^n^ represents RecB^n^, and B^h^CD stands for RecB^h^CD. The dissociation constant *K*_d_ is defined in the equation (1):

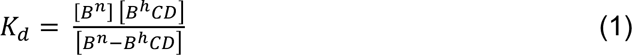

Since RecB^h^CD can only bind to the Ni-NTA beads when it forms a complex with the ^His^RecB^n^, the amount of RecB^h^CD eluted from the Ni-NTA beads equals the amount of RecB^h^CD-RecB^n^ complex formed. The ratio of [RecB^h^CD-RecB^n^] to total [RecB^n^] is given in equation (2):

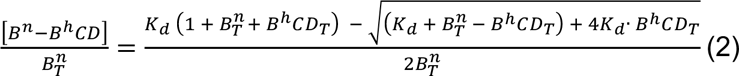

where B^n^_T_ and B^h^CD_T_ stand for the total concentrations of RecB^n^ and RecB^h^CD, respectively. Data were analyzed by nonlinear least square (NLLS) analysis using Scientist (Micromath, St. Louis, MO) and equations (1) and (2) to determine *K*_d_.

### χ-specific ssDNA formation assay

χ-specific ssDNA formation assay were performed by mixing 10 μM (nts) radioactively labeled dsDNA, 0.1 nM RecBCD (or 0.1 nM RecB^h^CD plus 1 μM RecB^n^) and 1 μM SSB in 25 mM Tris-acetate, pH 7.5, 10 mM Mg(OAc)_2_ unless noted otherwise. This mixture was incubated at 37°C for 3 min, and the reaction is initiated by addition of ATP to 5 mM. 10 μL aliquots were removed at each specified time points and quenched with 65 mM EDTA followed by deproteinization with 2% (w/v) SDS and 1 mg/ml proteinase K (Roche, Indianapolis, IN). These were loaded onto a 1% agarose gel and run at 40 V for 16 hr. The gel was dried and exposed to a phosphor screen and quantified on a Storm phosphorImager (GE Healthcare, Piscataway, NJ).

The fraction of χ ssDNA produced *f*_Chi_(*t*) at time *t* was calculated using equation (3):

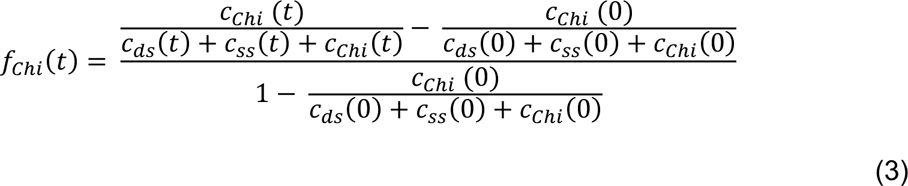

where *c*_Chi_(*t*), *c*_ds_(*t*) and *c*_ss_(*t*) are the counts of the bands corresponding to the χ ssDNA, dsDNA and ssDNA at time *t*, respectively. Since DNA is in excess, all the RecB^h^CD-RecB^n^ complexes should be bound to dsDNA. Thus, the fraction χ ssDNA produced should reflect the ratio of RecB^h^CD-RecB^n^ complexes formed to total [RecB^h^CD]. This relation can be expressed in terms of *K*_d_, total [RecB^n^] and [RecB^h^CD] as given in equation (4):

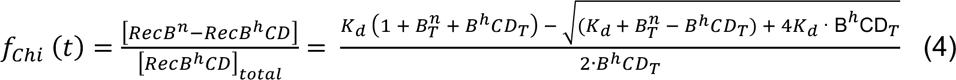

Data were analyzed using NLLS with Scientist (Micromath, MO) to determine *K*_d_.

### χ-dependent reversible inactivation assay

χ-dependent reversible inactivation of DNA unwinding was performed as described ^21^. The reaction mixture was the same as the regular χ ssDNA assay (25 mM Tris-acetate, 5 mM ATP, 1 μM SSB monomers) except that the concentrations of RecBCD and RecB^h^CD were 0.1 nM and 0.2 nM, respectively, and the concentration of Mg(OAc)_2_ was 1 mM. Assay temperature was at 37 °C, and 40 min after the reaction was initiated by addition of 5 mM ATP, excess Mg(OAc)_2_ was added to a final concentration of 10 mM. Aliquots were removed at specified time points and processed as described above.

### Joint molecule formation assay

Coupled joint molecule formation reactions for RecA-RecBCD, RecB^h^CD, and RecB^h^CD-RecB^n^ were conducted as described earlier ^7, 12^. Briefly, reactions were carried out in 25 mM Tris-acetate (pH 7.5), 8 mM Mg(OAc)_2_, 5 mM ATP, 1 mM dithiothreitol, 1 mM phosphoenolpyruvate, 2 units/ml pyruvate kinase, 10 μM (nucleotides) 5’-end-labeled NdeI-linearized χ^+^-pBR322 3F3H dsDNA, 20 μM (nucleotides) supercoiled circular χ^+^-pBR322 3F3H DNA, 5 μM RecA protein, 2 μM SSB protein and either 0.3 nM RecBCD, 0.9 nM RecB^h^CD, or 0.3 nM RecB^h^CD with 1 μM RecB^nuc^. When the RecB^n^ concentration was varied as indicated, reactions contained 0.3 nM RecB^h^CD. Reactions were terminated at indicated times by adding stop buffer. The reaction products were separated on a 1% (w/v) TAE (40 mM Tris-acetate (pH 8.2) and 1 mM EDTA) agarose gel at 600V·h, visualized, and quantified using Amersham Biosciences Storm 840 PhosphorImager and ImageQuant software.

### UV sensitivity test

*E. coli* strain AB1157 1−*recCBD::*FRT containing pMS421 (for *lacI^q^* expression) pPB520 (RecC expression), pPB800 (RecB and RecD expression), pPB800-TAAs (for RecB^927^ expression without RecB^n^ and RecD expression), and pGB^n^ (RecB^n^ expression on a low copy number plasmid pGL10) were gown overnight in LB broth containing spectinomycin (20 µg/ml), chloramphenicol (25 µg/ml), ampicillin (100 µg/ml) and kanamycin (30 µg/ml) antibiotics. An aliquot (50 μl) of an overnight culture was inoculated into 5 ml of LB medium and was allowed to grow till early log phase (OD_600_ = 0.5) at 37 °C. Serial dilutions of cells were made, and 5 μl of diluted cells were spotted on LB agar plates. The plates were irradiated with UV light (254 nm) using a Spectroline short wave UV lamp (model-XX-15F, Spectronics corporation, NY, USA) for 0, 4, 8 and 12 seconds at a UV dose of 0.3 J/m^2^/s. The colonies that survived UV irradiation after incubation at 37 °C for 20 hours in the dark were scored.

### T4 *2*^−^ plaque assay

*E. coli* strain AB1157 1−*recCBD::FRT* expressing RecBCD mutant enzymes from plasmids pPB520, pPB800, pPB800-TAAs, and pGB^n^ were grown overnight in LB broth containing chloramphenicol (25 µg/ml), ampicillin (100 µg/ml) and kanamycin (30 µg/ml) as needed. The pPB800, pPB800-TAAs, pPB520, and pGB^n^ are the low-copy expression vectors having 10-20 copies per cell and these proteins were expressed under LacI^q^ repression (from the pMS421 plasmid) without IPTG induction. An aliquot (50 μl) of an overnight culture was inoculated into 5 ml of LB medium containing suitable antibiotics, was allowed to grow till early log phase (OD_600_ = 0.5) at 37 °C. Early log cell culture (0.1 ml) was mixed with 3 ml of LB top agar and spread on LB bottom agar plates. Plates were allowed to dry for 15 min. Serial dilutions of the phage stock (10^9^ plaque forming units/ml) were made in suspension medium (50 mM Tris-HCl, pH 7.5, 100 mM NaCl, 1 mM MgSO_4_, 0.01% gelatin), and 5 μl of each dilution was spotted on LB agar plate containing bacterial cells. These plates were incubated at 37 °C overnight, and the number of countable plaques formed under lower dilution was counted. T4 or T4 2− phage titers from strains expressing RecBCD and derivatives were divided by the titer from strain AB1157 *1−recCBD::*FRT with an efficiency of plating (EOP) about 3×10^10^/ml.

### Lambda recombination and χ activity assay

The bacteriophage λ crosses were performed between λ phage 1081, λ phage 1082, λ phage 1083, and λ phage 1084 (described in experimental model and subject details section). Cross 1 was performed between phage 1081 × 1082, and cross 2 between phage 1083 × 1084. The frequencies of J^+^R^+^ recombinants of the mixed λ phage crosses (cross 1 and cross 2) from the *recBCD* null strains expressing RecBCD mutant enzymes from pPB520, pPB800, pPB800-TAAs, and pGB^n^ vectors determined by plating them on *E. coli* strain 594 (*sup*^0^) for recombinants, also on strain C600 (*supE*) for the total phage titer. The χ activity in these crosses was determined by the method of Stahl and Stahl ^22^ using the equation, 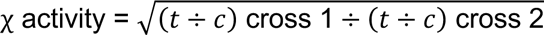, where (t ÷ c) is the ratio of turbid (t) to clear (c) plaques from cross 1 or cross 2, among J^+^R^+^ recombinants as described ^28^.

### Quantification and Statistical analysis

Data are presented as mean, ± Standard Error of the Mean (SEM) or Standard Deviation (SD), unless otherwise stated. At least three independent experiments have been performed for each experiment. Data from the protein-protein interactions using Ni-NTA magnetic beads and the χ-specific ssDNA formation assays were analyzed by nonlinear least square (NLLS) analysis using Scientist software (Micromath, MO). The gels were quantified using Gel Pro Analyzer (Media Cybernetics, Bethesda, MD). Data from the χ-dependent reversible inactivation, and the joint molecule formation assays were quantified using Amersham Biosciences Storm 840 PhosphorImager and analyzed using the ImageQuant software (GE Healthcare, Piscataway, NJ). Graphs and image processing were done using the Prism 5.0 (Graphpad, USA) and ImageJ (nih.gov) software programs.

